# Cultures of *Tyrophagus putrescentiae* experimentally infected with *Cardinium* and *Wolbachia* presented reduced fitness

**DOI:** 10.1101/2025.04.09.647973

**Authors:** Jan Hubert, Eliza Glowska-Patyniak, Stano Pekar

**Author notes:** Corresponding author:*E-mail address* (J. Hubert).

## Abstract

*Tyrophagus putrescentiae* is a cosmopolitan pest of stored food and animal feed. Mite populations differ in their microbiome composition, resulting in variability in their fitness. Cultures of the stored-product mite *T. putrescentiae* are often singly infected by one of intracellular bacterial genera *Cardinium* and *Wolbachia*. No naturally occurring multi-infected (*Cardinium/Wolbachia*) cultures have been observed.

Under laboratory conditions, we mixed two singly infected mite cultures, i.e., *Cardinium*-infected (5L and 5S) and *Wolbachia*-infected (5N and 5P) cultures, to obtain four experimental cultures (5LN, 5LP, 5SN and 5SP), each infected with each of the endosymbionts. Population growth was used as a fitness indicator, and PCR with specific primers for a single mite was used to identify the prevalence of *Cardinium* and *Wolbachia* in the parental and mixed cultures.

The population growth of the experimental cultures was lower than that of the parents at the beginning of the experiment. After six months, 5SN and 5SP exhibited greater fitness than 5LP and 5LN. The population growth of the 5SN and 5SP cultures did not differ from that of the parental cultures 5P and 5S. *Cardinium*-infected mites prevailed in 6 cultures, *Wolbachia*-infected mites in 5 cultures, and asymbiotic individuals in 1 culture after six months. The proportion of *Cardinium-*infected individuals (45%) was greater than that of *Wolbachia*-infected individuals (25%), indicating that *Cardinium* had the ability to overcome *Wolbachia*. Although the cultures were multi-infected, doubly infected mites were rare (7%).

These results suggest that *Cardinium* is responsible for cytoplasmic incompatibility in *T. putrescentiae*. The results revealed that the presence of the intracellular symbionts *Cardinium* and *Wolbachia* strongly influenced mite population growth in the experiments. These data support the importance of the microbiome in this pest.

## 1. Introduction

The stored-product mite *Tyrophagus putrescentiae* (Schrank, 1781) is an important pest found in stored products, such as cereals, dried ham and sausages, cheeses, and dog food (Robertson, 1961; Stejskal et al., 2015; Zhang et al., 2018; Olivry and Mueller, 2019). The ingestion of food contaminated by mites at high density can induce anaphylactic shock (Sanchez-Borges et al., 2005). Analyses of the stored grain revealed that a majority of the samples, i.e., 14% of the samples (N=514), had a lower population density than 1 ind.g^−1^ grain; however, 1% of the samples were infested with 5 or more individuals g^−1^ grain (Stejskal and Hubert, 2008). However, samples with low mite population density can reach massive biomass under suitable conditions (Aspaly et al., 2007). An effort to establish a model of population growth on the basis of a range of environmental conditions, both physical and biological, is essential for providing early warning of mite infestations (Sanchez-Ramos and Castanera, 2001, 2005; Pekar and Zdarkova, 2004; Collins, 2012); however, biotic factors are usually omitted because of a lack of data. Recent studies have shown that *T. putrescentiae* is colonized by a diverse microbiome and that different cultures have various fitness traits (Erban et al., 2016; Hubert et al., 2021a).

Many arthropods and nematodes are hosts of intracellular bacteria, such as *Wolbachia* and *Cardinium* (Breeuwer et al., 2012; Schneider et al., 2012). These bacteria are maternally transmitted and can manipulate host reproduction and mating behavior (Vala et al., 2004) to achieve successful distribution in host populations (Perlmutter and Bordenstein, 2020). The reproductive manipulations include male killing, feminization, thelytokous parthenogenesis, and cytoplasmic incompatibility (CI), all of which lead to an increased proportion of infected females in the host population (Perlman et al., 2008; Ros and Breeuwer, 2009). However, in some systems, symbionts can also be mutualistic and provide nutrition or other benefits to their host (Nikoh et al., 2014).

To our knowledge, the microbiomes of 10 cultures of the stored-product mite *Tyrophagus putrescentiae* (Schrank, 1781) have been identified. Among them, 3 (e.g., the 5L and 5S cultures, Chinese) are singly infected by *Cardinium,* 2 are singly infected by *Wolbachia* (i.e., the 5N and 5P cultures), and the remaining 5 do not contain intracellular bacteria (Erban et al., 2016; Lee et al., 2019; Hubert et al., 2021a; Xiong et al., 2023; Klimov et al., 2024). No naturally occurring multi-infected *Cardinium/Wolbachia T. putrescentiae* cultures are known. The genomes of *Wolbachia* from *T. putrescentiae* (wTPut) were 0.9 and 1 Mb in size (GenBank accession Nos. GIJY0 and JAUEMM01), respectively, and formed a new lineage with the phylogenetically unrelated and ecologically dissimilar gall mite *Fragariocoptes setiger* (NZ_JACVWV01) (Klimov et al., 2022) classified as supergroup Q (Glowska et al., 2015; Klimov et al., 2024; Hubert et al., 2025b). The results of phylogenetic analyses of *Cardinium* from *T. putrescentiae* (cTPut) (JANAVR01 and JAZHET01) genomes differed from those of *Cardinium* from other mites, i.e.*, Cardinium* from *Brevipalpus califormicus* (cBcaIN1: JAACGE01, cBcaIN2: JAACGD01), *Brevipalpus yothersi* (cByotN1: JAACGG01) and *Dermatophagoides farinae* (CP101107-9 and VMBH01) (Xiong et al., 2023; Hubert et al., 2025a). The cTPut genome is more similar to that of the *Cardinium* strain from the insect *Sogatella furcifera* (CP022339) (Zeng et al., 2018). cTPut has a smaller genome (1.05 Mb) than *Cardinium* from *D. farinae* (cDFar) (Xiong et al., 2023). The cDFar genome contains two plasmids, which are absent in cTPut; cDFar is found only in females, whereas cTPut occurs in both sexes of *T. putrescentiae* (Erban et al., 2020; Hubert et al., 2021b). The presence of wTPut in males/females is not clear. Both cultures 5N and 5P are inhabited by the same genotype of the wTPut Q group (Glowska et al., 2015; Klimov et al., 2024; Hubert et al., 2025b). Similarly, cTPut exhibits the same genotype in the 5S and 5L cultures (Hubert et al., 2025a). Both cTPut and wTPut are transmitted maternally, as indicated by the microbial profile of *T. putrescentiae* eggs (Hubert et al., 2021a). It is currently unknown whether CI exists for *T. putrescentiae*.

Mites respond to suitable conditions by accelerating their population growth, and this property is used in feeding biotests (Matsumoto, 1965). These biotests start from low but known numbers of mites, and the final density of nymphs/adults is compared after the observation period, usually 21 or 28 days (Hubert et al., 2005; Erban et al., 2015). Population growth has been recalculated as the intrinsic growth rate in some studies (e.g., Aspaly et al. (2007)).

The life cycle of *T. putrescentiae* from egg to adult lasts 9.4 days under optimal conditions (Sanchez-Ramos and Castanera, 2001). Previous experiments have revealed that a mixed culture (5LN) established from *Cardinium* (5L)- and *Wolbachia* (5N)-infected mites presented lower population growth than did parental singly infected cultures after 21 days (Hubert et al., 2021b). In population-level samples, *Cardinium* reduced *Wolbachia* abundance 3-fold, based on barcode sequencing, and 10-fold, based on qPCR (Hubert et al., 2021b). It is currently unknown whether CI exists for intracellular bacterial infections of *T. putrescentiae*. However, the decrease in the population of the mixed culture (5LN) in the experiment may have been caused by CI.

In the present study, we sought to compare population growth in experimental cultures in relation to the prevalence of intracellular symbionts. The experiment was based on previous studies performed in the same system (Hubert et al., 2021b), i.e., the mixing of individuals from parental cultures the comparison of symbiont prevalence and host fitness in parental and mixed cultures. The modifications were as follows: (i) more parental cultures were infected with *Cardinium* (5L and 5S) and *Wolbachia* (5N and 5P), resulting in four experimental cultures (5LN, 5LP. 5SN and 5SP); (ii) The cultures were sampled 3 times during five or six months of experimental growth, which was repeated in three separate experiments. The observed parameters were population growth and bacterial prevalence in the mites measured by PCR with taxon-specific primers.

## 2. Materials and methods

### 2.1. Parental cultures

For the experiments, four cultures of *T. putrescentiae* that were infected with either *Cardinium* (5L and 5S) or *Wolbachia* (5N and 5P) were used (Table 1). The cultures were maintained at the Czech Agricultural Research Center (CARC; Crop Research Institute until 2024), Prague, Czechia. The mites were kept in 70 mL IWAKI tissue culture flasks with a surface area of 25 cm^2^. These flasks were placed in Secador desiccators by Bel-Art Products, which maintained a relative humidity of 85% through a saturated KCl solution. The desiccators were kept in darkness under controlled humidity (75% RH) and temperature (25 ± 1 °C). The mites were fed a diet consisting of wheat germ and Mauripan-dried yeast extract (*Saccharomyces cerevisiae*) at a 10:1 w/w ratio. The diet was mill-powdered, sieved (mesh size, 500 µm), and heated to 70 °C for 0.5 h before being fed to the mites (Erban and Hubert, 2008). Every culture was renewed monthly by transferring approximately 5,000 live mites from the cap or surface of the flask into a new flask containing 0.3 g of SPMd. In our experience, adult mites usually escape from the diet and aggregate on the plugs and flask surface, whereas juveniles and eggs stay on the diet at the bottom of the flask. Adult mites can be easily sampled from plugs and surfaces. The cultures were sampled at the time of the first manipulation experiment. A previous analysis revealed that intracellular symbiont infection is stable over time (Erban et al., 2016; Hubert et al., 2021b). We ran two independent samplings after 5 months for symbiont prevalence determination.

**Table 1.** List of singly infected cultures of *Tyrophagus putrescentiae* used in this study. The origin of the culture, symbionts and rearing diet are provided. The numbers of males and females in the culture originated from a previous study (Erban et al. 2016) based on the inspection of mites in lactic acid under a compound microscope. The symbiont profile of the mite microbiome represented as the mean (±standard deviation) values from previously published data (Hubert et al. 2021a).

| ID | Population | Collector | Year | Diet | Site | Male | Females | Symbionts |
| --- | --- | --- | --- | --- | --- | --- | --- | --- |
| 5L | laboratory | E. Zdarkova | 1996 | SPMd | grain, Bustehrad, Czechia | 55 | 63 | *Cardinium* 45±3 % |
| 5N | dog | J. Hubert | 2007 | F | food-producing factory, St. Louis, Missouri, USA | 32 | 109 | *Wolbachia* 62±6 % |
| 5P | Phillips | T. W. Phillips | 2014 | SPMd | laboratory strain, Manhattan, Kansas, USA | 96 | 157 | *Wolbachia* 60±2 % |
| 5S | ham | A. Sala | 2013 | SPMd | food-producing factory, Cesena, Italy | 181 | 170 | *Cardinium* 53±4 % |

**Legend:** SPMd – stored-product mite diet, F – dog kernel, Purina Pro Plan FOCUS Adult Sensitive Skin & Stomach Salmon & Rice Formula Dry Dog Food.

### 2.2. Experimental cultures

Sex determination is not possible for live *T. putrescentiae* mites. To create mixed cultures, we transferred 10 unsexed adults from a *Cardinium*-infected culture (5L or 5S) and another 10 from a *Wolbachia*-infected culture (5N or 5P) into a fresh flask with fresh diet. The following combinations were obtained: 5LN, 5LP, 5Sm and 5SP. Each flask contained 20 mites, and 6 replicates were used for each combination (Fig. 1). Each replicate was examined in a separate flask that contained 0.3 g of diet. This experiment was repeated three times after half a year.

**Fig. 1.**
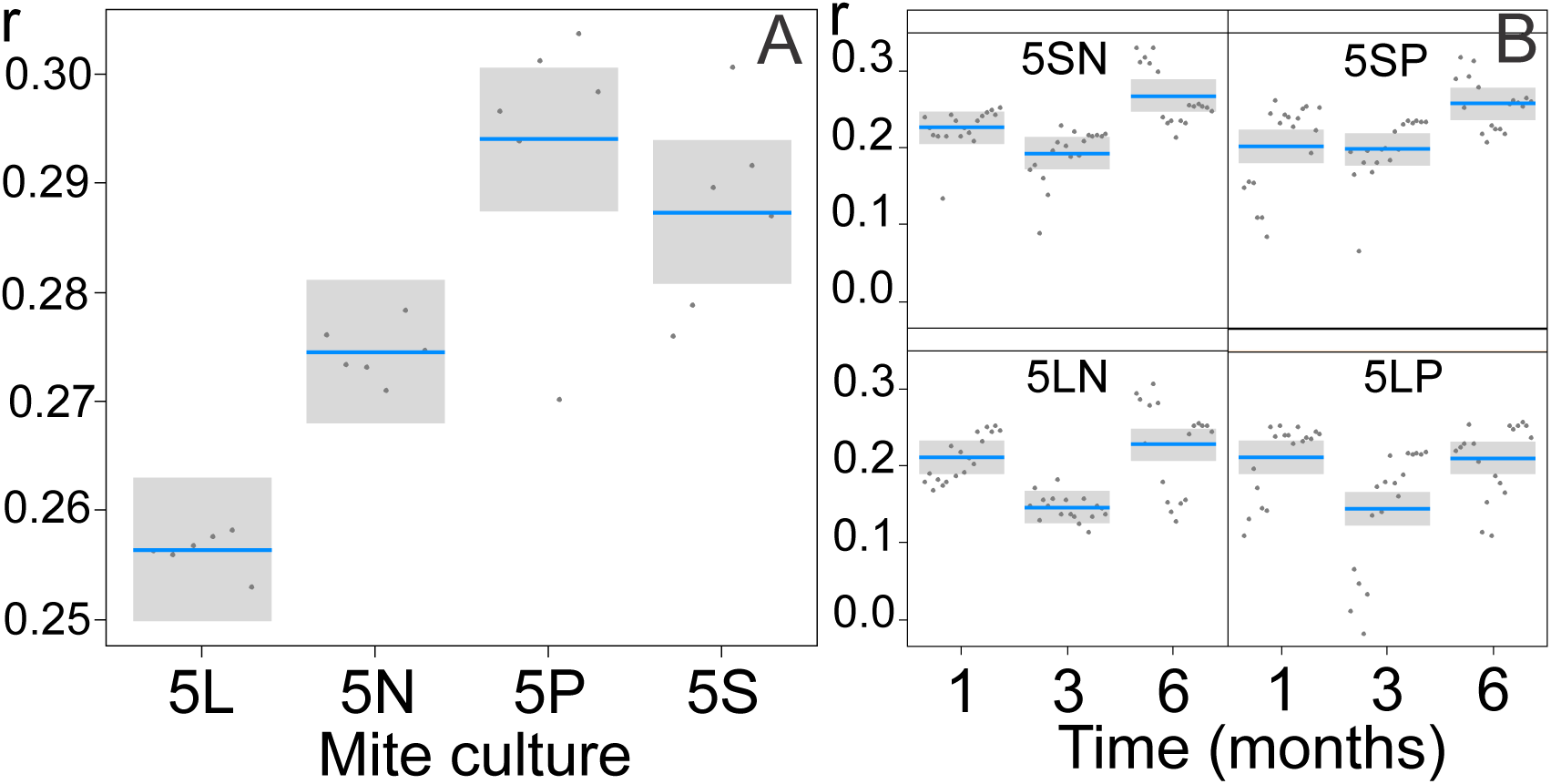
The experimental design was as follows: the mixing of individuals from two *Cardinium* (5L and 5S) and two *Wolbachia* (5N and 5P) singly infected cultures resulted in four experimental cultures (5LN, 5LP, 5SN and 5SP).

The flasks were stored separately in desiccators for culture to avoid cross contamination by mites. Every culture was renewed in the same way as the parental cultures, and the mite adults were collected and used for symbiont prevalence determination (after 2, 3, 4 and 5/6 months) and population growth tests (after 1, 3 and 5 or 6 months). The reason for the different durations is that experiment Nr. I. was terminated after 5 months because *Cardinium* eliminated *Wolbachia*. However, no elimination was observed after 5 months in experiments Nr. II and Nr. III, and these experiments were terminated after 6 months.

### 2.3. DNA extraction

For DNA extraction, the mites were removed from the plugs and stored in 80% ethanol and transferred to sterile petri dish.. We pooled all mites from the six replicated chambers, and 30 mites were selected randomly and used for DNA isolation separately. The exceptions were the parental cultures, wherein 30 mites from the beginning of the experiment and 30 mites after 5 months from preparental cultures were analyzed. For every samples, single mite was removed from ethanol and transferred into a 0.2 mL thin-walled tube (Thermo Scientific™, cat no: AB0620) that contained 25 μL of DEPL25 STARTLBlue reagent (cat no: D226) for alkaline lysis, and the mixture was blue. The tube was then heated to 95 °C for 20 minutes using a C1000 Thermal Cycler (Bio-Rad, Hercules, CA, USA). After heating, the tube was cooled to room temperature, and 25 μL of DEP-25 STOP solution was added and mixed by vortexing, after which the solution became transparent. The isolated DNA was immediately subjected to PCR analysis.

### 2.4. Intracellular bacterial prevalence determination via PCR

The analyses were based on 3 PCRs for every isolated DNA sample (1 mite). PCRs were carried out using EmeraldAmp Master Mix (catalog number: RR310A, Takara Bio). The master mix contained an optimized buffer, a polymerase, a dNTP mixture, gel loading dye (green), and a density reagent in a 2X premix format. Subsequently, ddH_2_O and primers were added to the mix. The amplification process was carried out via a C1000 Thermal Cycler (Bio-Rad, Hercules, CA, USA).

i. For *Wolbachia* detection, the WpF (5’-TTGTAGCCTGCTATGGTA-3’) and WpR (5’-GAATAGGTATGATTTTCA-3’) primers (O’Neill et al., 1992) were used with the following amplification profile: initial denaturation at 94 °C for 5 minutes; 35 cycles of 95 °C for 60 seconds, 52 °C for 60 seconds, and 72 °C for 60 seconds; and a final extension at 72 °C for 5 minutes. The positive control contained DNA from 5N cultures (population-level sample), and the negative control contained ddH_2_O instead of the DNA sample.
ii. The Card4 (5’-CTTAACGCTAGAACTGCGA-3’) and Card6 (5’-TCAAGCTCTACCAACTCC-3’) primers (Kopecky et al., 2013) were used for *Cardinium* detection with the following protocol: initial denaturation at 94 °C for 5 minutes, followed by 35 cycles of denaturation at 94 °C for 50 seconds, annealing at 56 °C for 50 seconds, extension at 72 °C for 60 seconds, and a final extension at 72 °C for 10 minutes. The positive control contained DNA from 5L cultures (population-level sample), and the negative control contained ddH_2_O instead of the DNA sample.
iii. Universal bacterial primers (F27 and 1492R) (Lane, 1991) and a protocol described previously (Hubert et al., 2012) were used to identify the presence of 16S DNA in the samples. The positive control contained the 5L culture (population-level sample), and the negative control contained ddH_2_O instead of the DNA sample.

The PCR products were observed on a 1% gel using the GeneSnap system (Syngene InGenius LHR2 Gel Imaging System; cat. no: 316616). The size of the products was determined using a 50 bp ladder (Generuler 50bp, cat no: SM0373, Thermo Fisher Scientific). The amplification process was successful when the PCR products were visible and were of the expected size. The asymbiotic mite individuals were identified on the basis of the presence of the product from universal bacterial primers and the absence of the product from the *Cardinium* and/or *Wolbachia* primers.

### 2.5. Population growth test

Mites were collected from the plugs and surfaces of rearing flasks of parental or experimental cultures and then transferred to separate Petri dishes.

Ten randomly selected adult mites were transferred from a Petri dish to new flasks that contained 0.01±0.005 grams of SPMd. Mites cannot be sexed without being placed on slides, and sex identification via a compound microscope is not possible (Erban et al., 2016). The flasks were kept under the same conditions as those used for mite rearing. After 21 days, the experiment ended, and mites were counted using a dissection microscope (Hubert et al., 2016).

### 2.6. Statistical analysis

For the data analysis, we used a measure of population fitness, which was calculated as the intrinsic rate of population increase (*r*), assuming exponential population growth, as the time was short and resources were not limited. We used the following formula for density-independent continuous population growth: *N*_t_ = *N*_0_e*^rt^*, where *N_t_* is the final mite density, *N*_0_ is the initial mite density, and *t* = 21 days represents the duration of mite population growth in the experiment. The intrinsic rates had a distribution that did not differ from normal distribution; thus, differences among populations were studied using a general linear model (LM). To test for consistency (repeatability) in the rate in the same population mixtures grown in three experiments (i.e., Nr. I, II and III), we used linear mixed-effect models (LMEs) from the nlme package (Pinheiro and Bates, 2000). The intrinsic rate was the response variable, and population type and time were explanatory variables. The experiment had a random effect. We calculated the intraclass correlation coefficient (ICC) from the variance estimated as a measure of consistency. The prevalence of endosymbionts in singly infected populations was analyzed with generalized linear models with binomial error structure (GLM-b) (Pekar and Brabec, 2016). The prevalence rates (after applying an angular transformation to approach a normal distribution) in mixed populations were compared among population types and times using LMEs to estimate consistency in prevalence using ICC. Eventually, the effect of prevalence on population growth was studied using LM. Significant differences among factor levels were assessed by nonoverlap of 95% confidence intervals. All analyses were performed in R (R Development Core Team, 2023).

## 3. Results

### 3.1. Population growth

The intrinsic growth rates of singly infected cultures of *T. putrescentiae* (i.e., 5L, 5S, 5N, and 5P) significantly differed (LM, F_3,20_ = 27.6, P < 0.001; Fig. 2A). One culture of those infected with *Cardinium* (5S) and one of those infected with *Wolbachia* (5N) exhibited a significantly greater intrinsic growth rate than the other two (5L and 5P, respectively) (Table 2). The four experimental cultures (5LN, 5LP, 5SN, and 5SP) presented significantly different intrinsic growth rates (LME, F_6,202_ = 2.7, P = 0.015) (Table 3). After one month, all four cultures exhibited a similar rate (Fig. 2B), which was lower than that observed for singly infected cultures (P< 0.05). After three months, the intrinsic growth rate was lower, but only for 5LP and 5LN was the difference significant compared with that after one month. After five months, all the cultures grew at significantly different rates than they did after three months (P< 0.05). Compared with 5LP and 5LN, 5SN and 5SP presented greater intrinsic growth rates. These two cultures had an intrinsic growth rate that was not significantly different from that of singly infected cultures of 5P and 5S. The low intraclass correlation coefficient (0.19) shows that the three experiments had low consistency.

**Fig. 2.**
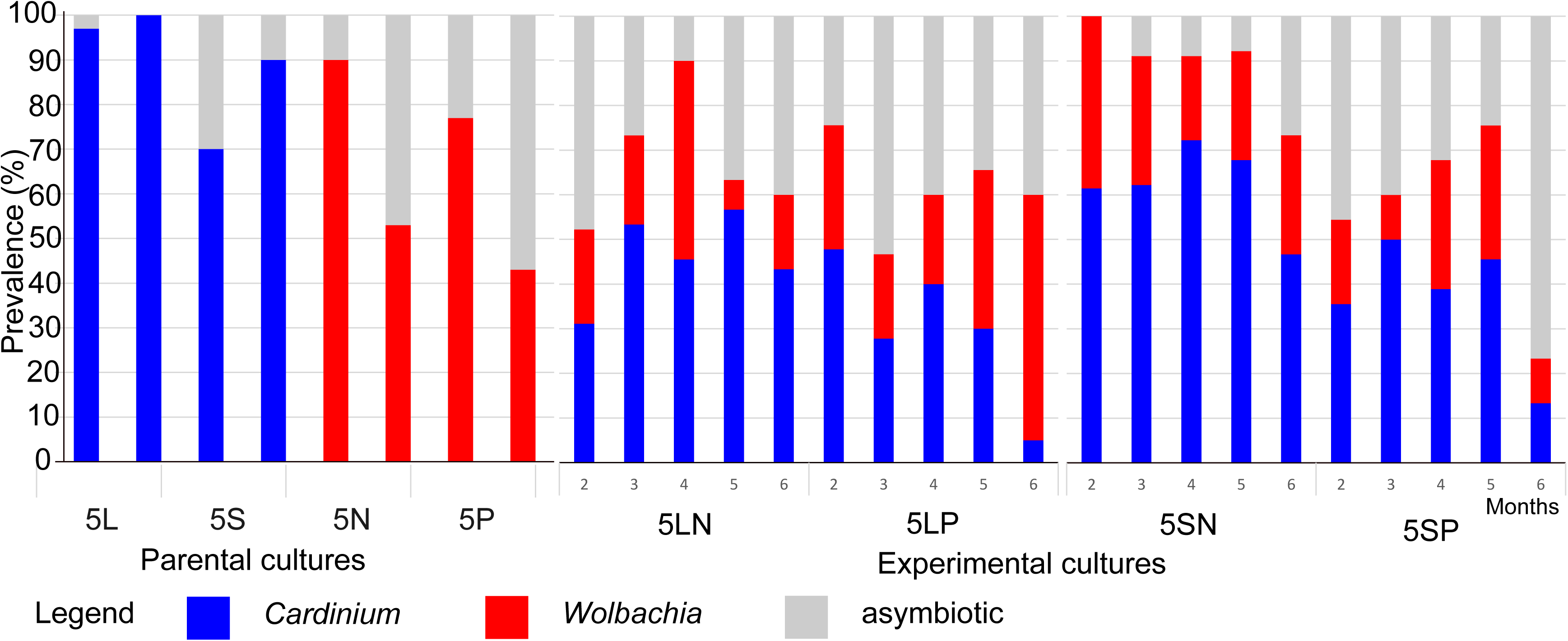
Comparison of the intrinsic rate of *Tyrophagus putrescentiae* population growth (*r*) among parental cultures (**A**) and among the four experimental cultures throughout the experiments (**B**). The horizontal lines represent the estimated means, and the gray boxes represent 95% confidence intervals.

**Table 2.**
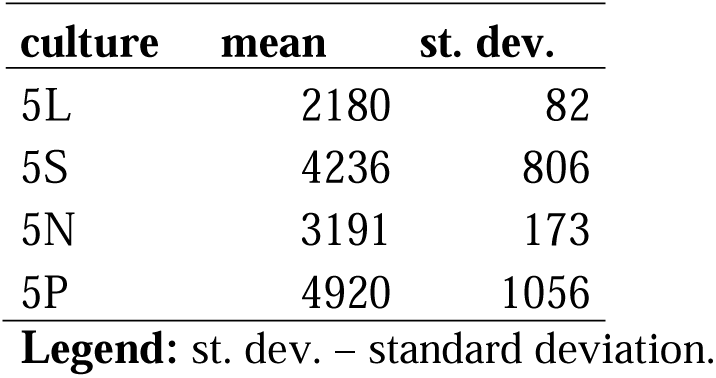
Results of the population growth test using parental *Tyrophagus putrescentiae* cultures. The population growth test was based on mite population growth for 21 days from the initial population of 10 individuals. The results of the test are expressed as the number of mite individuals, and the means and standard deviations are shown.

| culture | mean | st. dev. |
| --- | --- | --- |
| 5L | 2180 | 82 |
| 5S | 4236 | 806 |
| 5N | 3191 | 173 |
| 5P | 4920 | 1056 |

**Legend:** st. dev. – standard deviation.

**Table 3.** Results of the population growth experiment using experimental *Tyrophagus putrescentiae* cultures. Three separate experiments were performed, and after each period, the population growth tests were repeated. The population growth test was based on mite population growth for 21 days from the initial population of 10 individuals. The results of the tests are expressed as the number of mites per individual, and the means and standard deviations are shown.

| Culture | Time months | Experiment Nr.1 |  | Experiment Nr.2 |  | Experiment Nr.3 |  |
| --- | --- | --- | --- | --- | --- | --- | --- |
|  |  | mean | st. dev. | mean | st. dev. | mean | st. dev. |
| 5LN | 1 | 438 | 69 | 802 | 255 | 1750 | 258 |
|  | 3 | 255 | 73 | 233 | 119 | 197 | 58 |
|  | 5/6 | 3992 | 1682 | 255 | 96 | 1932 | 215 |
| 5LP | 1 | 277 | 191 | 1653 | 301 | 1585 | 236 |
|  | 3 | 47 | 63 | 433 | 238 | 882 | 175 |
|  | 5/6 | 1243 | 437 | 285 | 165 | 1930 | 254 |
| 5SN | 1 | 952 | 449 | 1170 | 315 | 1730 | 214 |
|  | 3 | 327 | 192 | 813 | 284 | 930 | 61 |
|  | 5/6 | 7987 | 2057 | 1325 | 223 | 2073 | 138 |
| 5SP | 1 | 170 | 93 | 1673 | 607 | 1548 | 607 |
|  | 3 | 401 | 217 | 658 | 221 | 1377 | 46 |
|  | 5/6 | 4975 | 2234 | 1040 | 161 | 2353 | 171 |

**Legend:** st. dev. – standard deviation.
**Remark:** The length of experiment Nr.1 was 5 months, whereas experiments Nr.2 and 3 had a duration of 6 months.

### 3.2. Prevalence of intracellular bacteria

The prevalence of intracellular bacteria in singly infected cultures (Table 4) differed significantly between cultures with different bacterial species (GLM, χ^2^_2_ = 89.4, P < 0.0001): the prevalence of *Cardinium* was consistently higher than the prevalence of *Wolbachia* in the parental cultures (Fig. 3A). However, the *Cardinium* prevalence in 5L was greater than that in 5S, and the *Wolbachia* prevalence in 5N was greater than that in 5P (χ^2^ = 50.0, P < 0.0001). There were differences among the experiments, and experiment Nr. 1 generally exhibited a higher prevalence than did experiment Nr. 2 (χ^2^_1_ = 14.3, P = 0.0002).

**Fig. 3.**
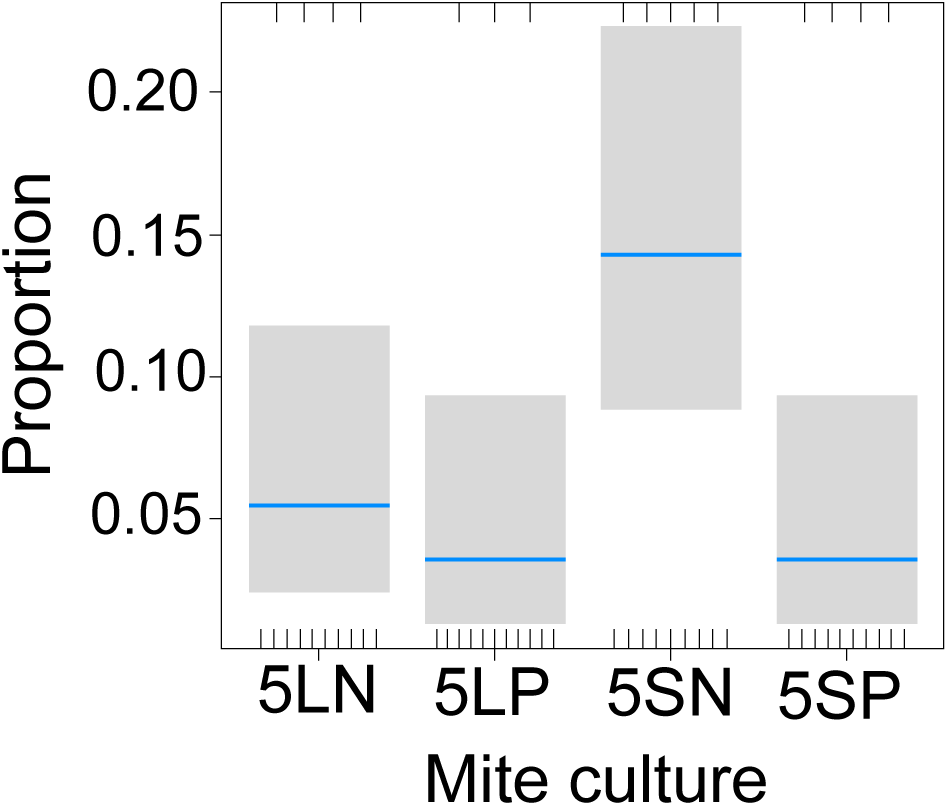
Comparison of the prevalence of *Cardinium* and *Wolbachia* in parental cultures of *Tyrophagus putrescentiae* (2 replicates) and in experimental cultures over six months.

**Table 4.** Numbers of mites infected by *Cardinium* or *Wolbachia* in parental cultures of *Tyrophagus putrescentiae* determined via PCR with specific primers. The single-mite PCR was performed with 60 replicates per culture. The sex ratio data were obtained from previous analyses (Erban et al. 2016). The bold symbol indicates that the proportion did not differ from the test ratio (single proportion test in PAST (Hammer et al. 2001)).

| Culture | Symbiont | Numbers of positive individuals |  |  | Test ratio 0.5 |  | male/<br>(male+female) | Test ratio males |  |
| --- | --- | --- | --- | --- | --- | --- | --- | --- | --- |
|  |  | P. | N. | R. | Z | P |  | Z | P |
| 5L | *Cardinium* | 59 | 1 | 0.98 | 7.436 | 0.001 | 0.45 | 8.251 | 0.001 |
| 5S |  | 48 | 12 | 0.80 | 4.647 | 0.001 | 0.5 | 4.647 | 0.001 |
| 5N | *Wolbachia* | 43 | 17 | 0.72 | 3.253 | 0.001 | 0.28 | **1** | **0** |
| 5P |  | 36 | 24 | 0.60 | **1.549** | **0.121** | 0.38 | **-0.32** | **0.75** |

Legend: P - number of positive PCR results, N – number of negative PCR results, R – P/N ratio.

In the experimental cultures, the prevalence of the two bacteria varied significantly among the cultures (LME, F_6,154_ = 2.42, P = 0.028). The total proportions of intracellular bacteria in all the experiments (Nr. I– III) at each time point tested were as follows: *Cardinium*, 45%; *Wolbachia*, 25%; asymbiotic, 37%. Approximately 7% of individuals were infected with both *Cardinium* and *Wolbachia* (N=1680). In all cultures, the prevalence of both bacteria fluctuated over six months but not significantly (LME, F_1,142_ = 0.3, P = 0.61). The consistency of the prevalence in the three experiments was nearly zero (ICC < 0.01).

After six months of experiments, *Cardinium*-infected individuals prevailed in 6 cultures, *Wolbachia* in 5 cultures and asymbiotic individuals in 1 culture (Table 5). In the 5LN culture, on average, the number of *Cardinium*-infected individuals was consistently higher than that of *Wolbachia*-infected individuals, the abundance of which decreased, so there were 2.6 times more *Cardinium*-infected individuals at the end. In the 5LP culture, on average, the number of *Cardinium*-infected individuals gradually decreased, so ultimately, there were 11 times more *Wolbachia*-infected individuals than *Cardinium-*infected individuals. In the 5SN culture, on average, the number of *Cardinium-*infected individuals was consistently higher than that of *Wolbachia*-infected individuals, which did not change; thus, in the end, there were almost 2 times more *Cardinium*-infected individuals. In the 5SP culture, on average, the numbers of both *Cardinium-* infected and *Wolbachia*-infected individuals decreased, so, in the end, a majority were asymbiotic individuals.

**Table 5.**
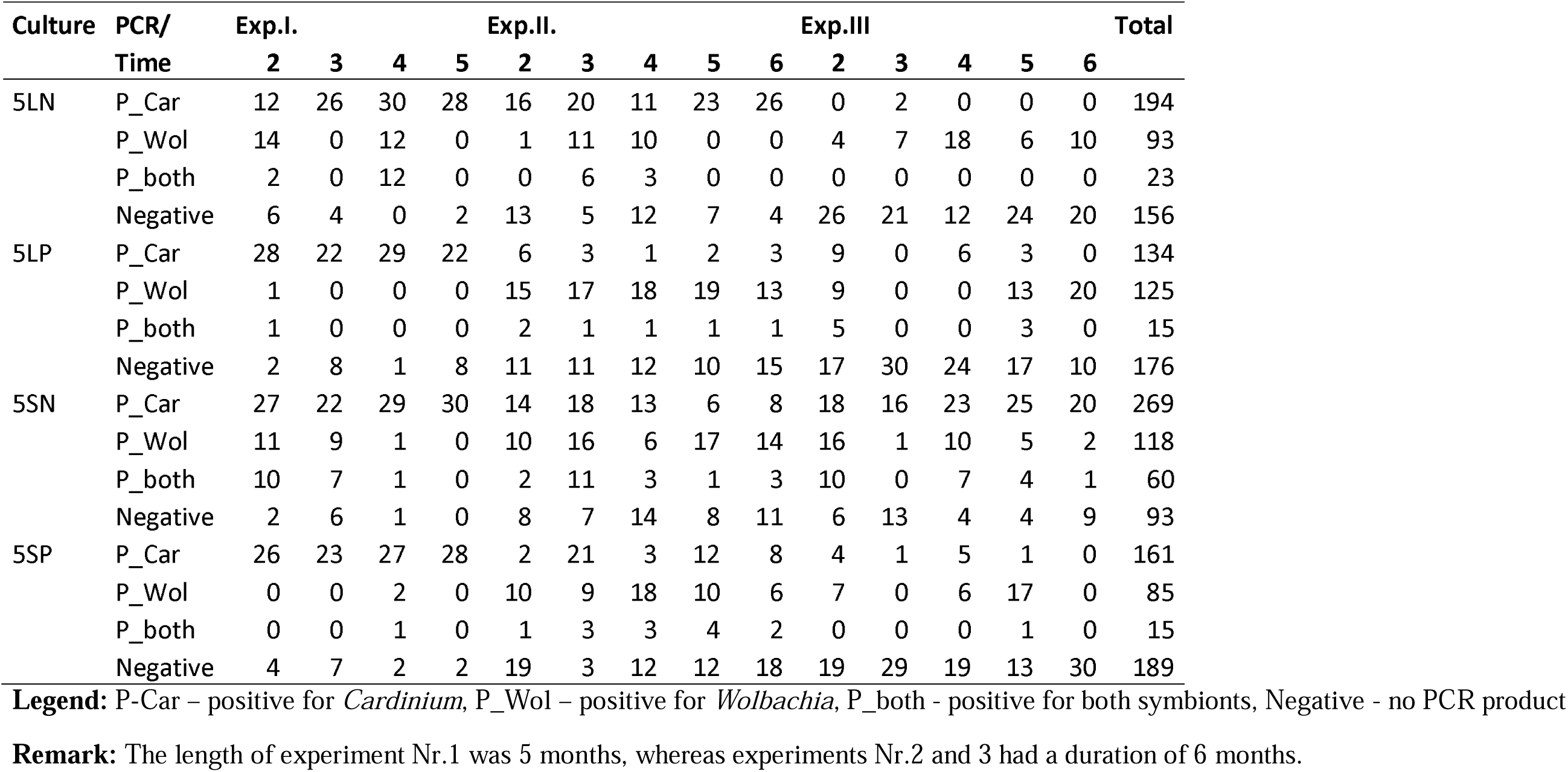
Numbers of mites infected by Cardinium or Wolbachia or doubly infected (Cardinium/Wolbachia) in the experimental cultures of Tyrophagus putrescentiae determined via PCR with specific primers. The single-mite PCR was performed in 30 replicates per culture, experiment and experimental period. The data were separated by month of culture growth and experiments.

Few individuals (7%, N=1680) in the experimental culture were doubly infected with both *Wolbachia* and *Cardinium*. The frequency of their occurrence did not change during the experimental period (GLM-qb, F_1,52_ = 13.0, P = 0.08) or in the experiment (GLM-qb, F_2,53_ = 4.0, P = 0.60), but with respect to culture type (GLM-qb, F_3,49_ = 47.1, P = 0.014), there were a significantly (3-fold) greater number of doubly infected individuals in 5SN than in the other cultures (Fig. 4).

**Fig. 4.**
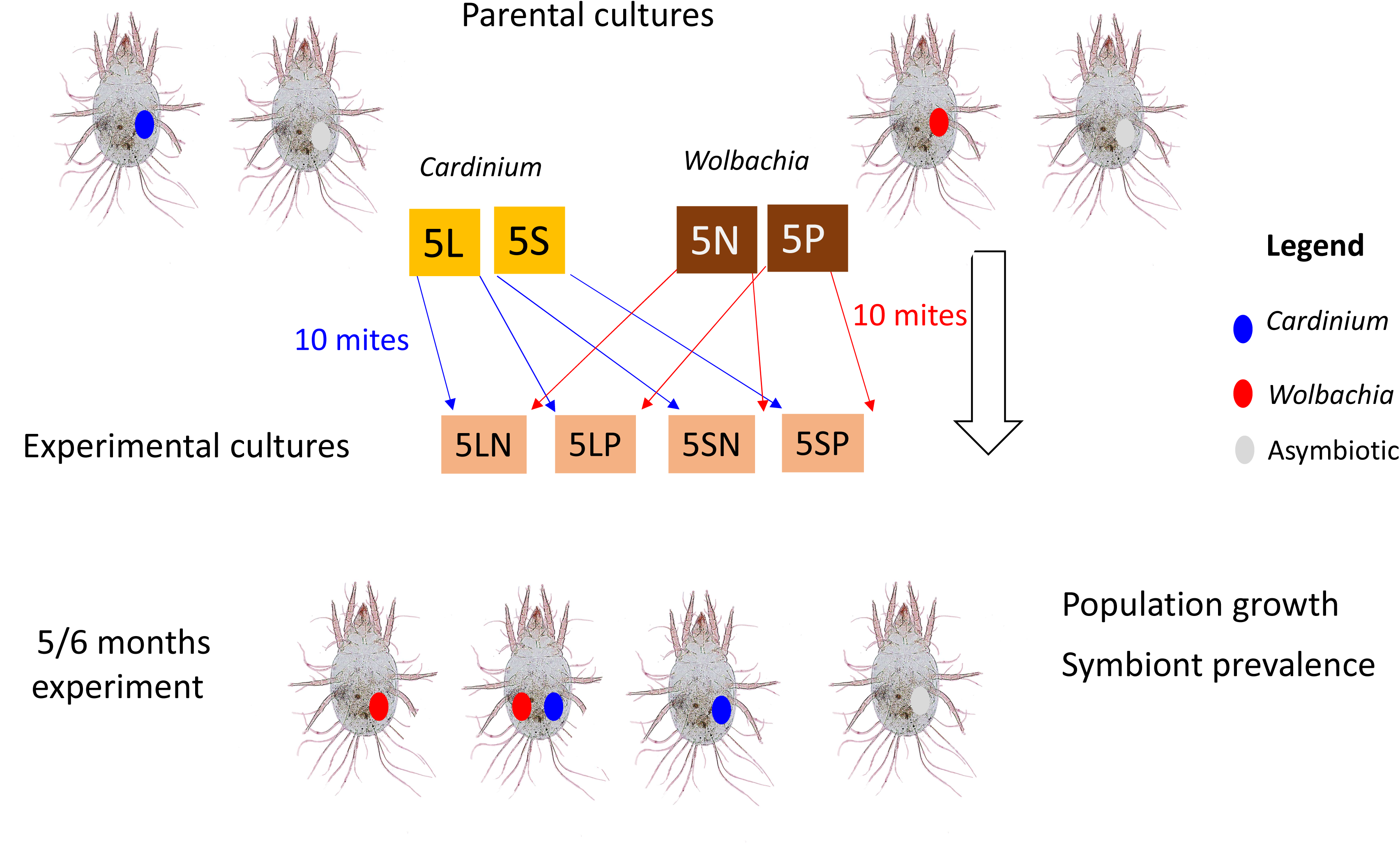
Comparison of the relative frequency of doubly infected individuals of *Tyrophagus putrescentiae* in the four mixed populations. The horizontal lines represent the estimated means, and the gray boxes represent 95% confidence intervals. The graph shows the proportions of doubly infected mites in the analyzed samples of experimental cultures during the whole experimental period.

By combining data on the intrinsic growth rate and prevalence, we found that the occurrence of *Wolbachia-*infected individuals (LM, F_1.34_ < 0.1, P = 0.97), *Cardinium*-infected individuals (LM, F_1.34_ = 1.4, P = 0.25) or asymptomatic individuals (LM, F_1.34_ = 2.9, P = 0.097) did not affect the intrinsic growth rate of the cultures after 21 days.

## 4. Discussion

This study revealed that, at the population level, multi-infected *T. putrescentiae* cultures are unstable and that the mixing of parental cultures resulted in a decrease in fitness after three months in experimental cultures. Only two experimental cultures achieved a similar population growth rate as the parental singly infected cultures did. We hypothesized that the process of infection is stochastic, i.e., it exhibits genetic drift (Jansen et al., 2008), and can result in all possible scenarios, i.e., *Cardinium* wins, *Wolbachia* wins, or, in some cases, neither wins and asymbiotic mites take over. Additionally, we showed that the prevalence of both symbionts could decrease, leading some individuals shifting to an asymbiotic state without intracellular symbionts. The mean generation time of *T. putrescentiae* was found to be 10 days under the optimal conditions (Sanchez-Ramos and Castanera, 2005), suggesting that in our study, there were approximately 12–15 generations. Previous analyses of artificial experimental cultures of *T. putrescentiae* have indicated that *Cardinium* wins over *Wolbachia* based on 21-day observations (Hubert et al., 2021b), i.e., on population-level samples of mites, *Cardinium* was able to reduce the abundance of *Wolbachia* 2.7-fold in the microbiome, as observed by sequencing profiling, or 10-fold, as observed by qPCR using taxon-specific primers (Hubert et al., 2021b). A previous study showed that *Cardinium* inhibits the growth of *Wolbachia* in experimental culture (5LN) after 21 days (Hubert et al., 2021b). Here, we obtained different results. In this study, *Cardinium*-infected individuals prevailed in 6 cultures, *Wolbachia-*infected individuals in 5 cultures and asymbiotic individuals in 1 culture after six months. However, the proportion of *Cardinium*-infected individuals (45%) was greater than that of *Wolbachia*-infected individuals (25%), indicating that *Cardinium* has the ability to overcome *Wolbachia* in *T. putrescentiae*.

The prevalence of *Cardinium* in doubly infected cultures of *T. putrescentiae* differs from that in cultures of *Tetranychus* spider mites (Zele et al., 2018). However, spider mites are infected by 3 intracellular symbionts, *Cardinium*, *Rickettsia*, and *Wolbachia* (Zele et al., 2018), while *Rickettsia* is not present in *T. putrescentiae* (Hubert et al., 2012, 2021a). The natural prevalence of symbionts in *Tetranychus* populations (N=16) decreased, with the greatest decrease observed for *Wolbachia* (61%), followed by *Cardinium* (12– 15%) and *Rickettsia* (0.9%–3%) (Zele et al., 2018). A laboratory experiment with spider mites revealed apparent loss of *Rickettsia* and *Cardinium*, but not *Wolbachia*, during 6 months of experiments (15 generations) (Zele et al., 2020). This finding indicates the stability of *Wolbachia* infection in *Tetranychus* populations, which is the opposite of the situation observed in *T. putrescentiae* in this study, where *Cardinium* was more stable.

The possible reasons for the changes in the proportions of intracellular parasites in the experimental cultures include CI and/or competition between bacteria through the manipulation of host pathways. It is currently unknown whether CI exists for *T. putrescentiae* infections. Both *Cardinium* and *Wolbachia* induce CI in *Tetranychus* and *Bryobia* mites (Gotoh et al., 1995, 2007; Breeuwer, 1997), although CI induced by *Wolbachia* differs among *Tetranychus* populations (Zele et al., 2020). In the spider mite *Panonychus ulmi*, *Cardinium* did not mediate sex allocation distortion in the experiments, and the effect of CI was not confirmed (Haghshenas-Gorgabi et al., 2023). In the doubly infected parasitoid wasp *Encarsia inaron*, *Wolbachia* caused CI, whereas *Cardinium* did not. In contrast, double infection of *Tetranychus urticae* with *Wolbachia* and *Cardinium* induced strong CI (Xie et al., 2016). The last possibility is bidirectional CI (Roed and Engelstadter, 2022), although it is usually associated with multiple *Wolbachia* lineages infecting the same host (Bordenstein and Werren, 2007). The presence of symbionts in males modifies sperm, leading to embryonic mortality in crosses with symbiont-free females as a condition for CI (Martinez et al., 2021). Previous observations have revealed significant differences in the known ratio of males to males+females (Erban et al., 2016), and the female proportion corresponded to the known proportion of *Wolbachia*-infected mites (Table 4). The interpretation of this finding is that *Wolbachia* infected females but not males of *T. putrescentiae*. However, this sex ratio was not observed in the present study. This finding led to the hypothesis that *Cardinium* caused CI and decreased the fitness of the experimental cultures. However, this finding needs to be verified in future studies.

The possible competition between *Cardinium* and *Wolbachia* was observed in doubly infected *Tetranychus piercei* (Zhu et al., 2012). The low proportion of doubly infected individuals in manipulated *T. putrescentiae* cultures should be the result of such competition. The indirect evidence of competition is the decrease in the intrinsic growth rate of all multi-infected cultures compared with parental cultures, which still occurred in some cultures during the experiment.

Our data support findings showing that microbiomes influence the fitness of the host (Gould et al., 2018). This is an important factor in the modeling of population growth for this pest. Mite population dynamics exhibit metapopulation patterns (Athanassiou et al., 2002; Stejskal, 2002; Hubert et al., 2006), and here, we found that the physiological features of the mites were determined by intracellular factors, resulting in a high degree of variability during the growth of the metapopulations.

## Author contributions

**Jan Hubert:** Writing, Designing and organizing experiments. **Eliza Glowska-Patyniak:** Writing, Designing and organizing experiments. **Stano Pekar:** Writing, Statistical analysis.

## Acknowledgments

The authors are grateful to two anonymous referees for valuable comments and to Eliska Tresnakova, Marta Nesvorna, and Martin Markovic for technical assistance.

